# Modelling the active SARS-CoV-2 helicase complex as a basis for structure-based inhibitor design

**DOI:** 10.1101/2020.11.03.366609

**Authors:** Dénes Berta, Magd Badaoui, Pedro J. Buigues, Sam Alexander Martino, Andrei V. Pisliakov, Nadia Elghobashi-Meinhardt, Geoff Wells, Sarah A. Harris, Elisa Frezza, Edina Rosta

## Abstract

Having claimed over 1 million lives worldwide to date, the ongoing COVID-19 pandemic has created one of the biggest challenges to develop an effective drug to treat infected patients. Among all the proteins expressed by the virus, RNA helicase is a fundamental protein for viral replication, and it is highly conserved among the *coronaviridae* family. To date, there is no high-resolution structure of helicase bound with ATP and RNA. We present here structural insights and molecular dynamics (MD) simulation results of the SARS-CoV-2 RNA helicase both in its apo form and in complex with its natural substrates. Our structural information of the catalytically competent helicase complex provides valuable insights for the mechanism and function of this enzyme at the atomic level, a key to develop specific inhibitors for this potential COVID-19 drug target.

## INTRODUCTION

Only a few approved drugs currently repurposed for treating COVID-19, a disease caused by the human coronavirus SARS-CoV-2, despite that its close relatives, SARS-CoV and MERS-CoV are responsible for multiple outbreaks earlier this century. As of September 2020, Remdesivir and Dexamethasone are used in clinical practice.^1,2^ Therefore, drugs need to be developed that can be used against the viral replication to help patients overcome the disease in the most severe cases.

Here we focus on determining the catalytically active complex structures of the SARS-CoV-2 RNA helicase, labelled as Non-structural Protein (NSP) 13 (Fig. 1). This protein is part of the Orf1ab polyprotein, which gets spliced to produce the enzymes required for viral replication. The RNA helicase performs two essential functions for the viral replication making it an ideal drug target. It is thought to perform the first step in the 5’-capping of the viral RNA by its triphosphatase function hydrolyzing the 5’-triphosphate group to form diphosphate-RNA.^3,4^ Furthermore, its main helicase function is to enable RNA translocation and unwinding in an ATP-dependent mechanism during viral replication.

**Figure 1.**
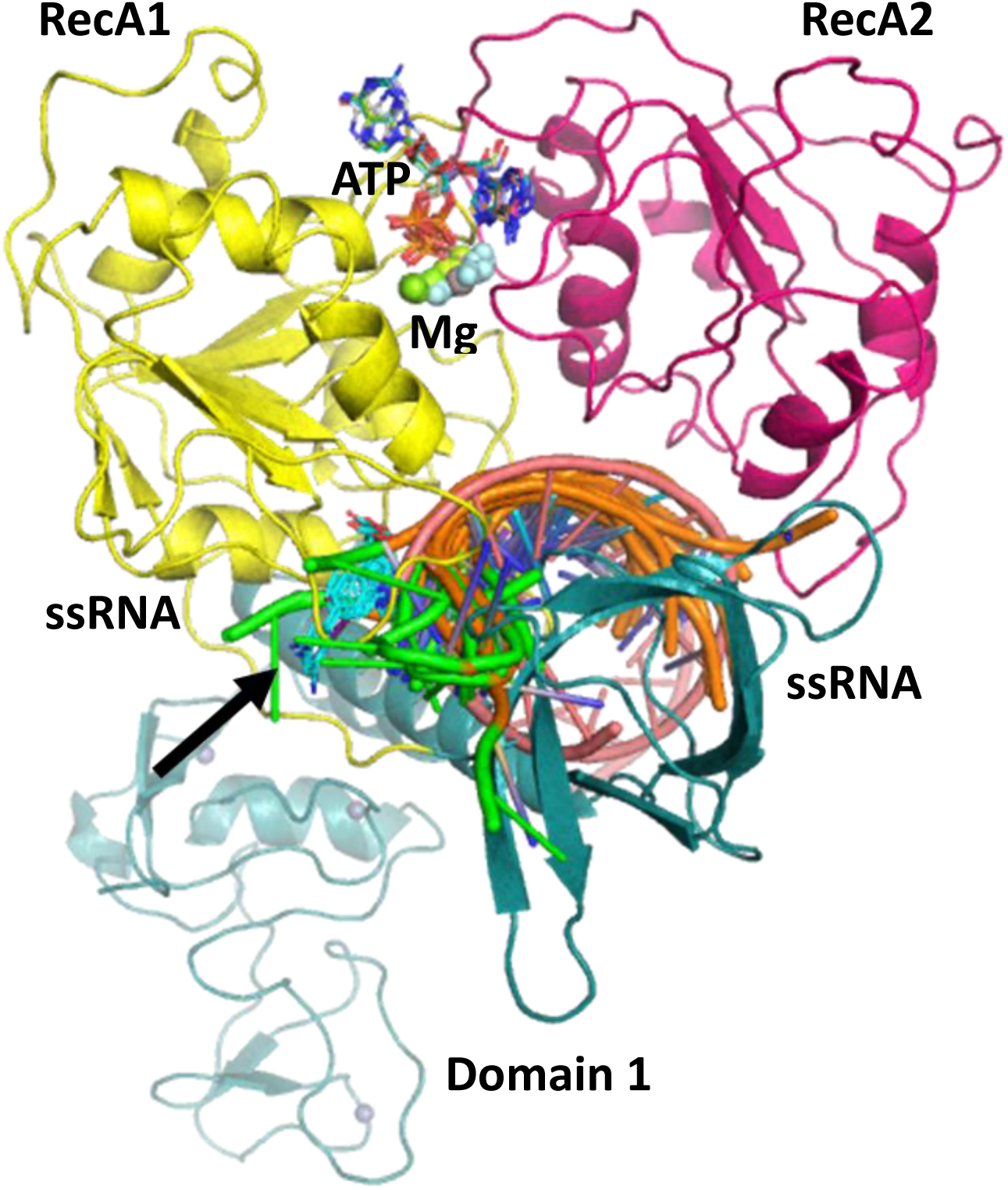
Cartoon representation of the RNA helicase NSP13 of SARS-CoV-2 monomer model composed of three domains: RecA1 (yellow), RecA2 (magenta), and Domain 1 (aquamarine). ATP analogues (sticks) along with Mg (green sphere) and single stranded nucleic acids are depicted from aligned homologous structures (complete list in the SI). 3’ ends of the nucleic acids present the same orientation in all chains (highlighted in green). Specific helicase inhibitor binding region with allosteric inhibitors displayed in cyan (black arrow).

Accordingly, numerous studies have already demonstrated that it is possible to develop potent inhibitors of viral helicases as antiviral agents.^5^

Coronaviral RNA helicases share high similarity. 600 out of the 601 residues of the SARS-CoV2 RNA helicase are identical to those of the SARS-CoV virus, and 70% match that of the MERS-CoV NSP13, demonstrating that these proteins are highly conserved within the coronaviridae family. Despite the importance of this target protein, currently only the apo structure is available crystallographically. There are no structures currently known of the RNA helicase bound with either inhibitors or nucleic acids, severely limiting the structure-based mechanistic understanding and the design of more potent drugs. Currently the only experimentally available information on SARS-CoV-2 helicase comes from a recent study by Shi et al., that offer a low resolution cryo-EM structure with ATP bound in the active site.^6^ They fit the crystal structure of the APO helicase from 6jyt to their cryo-EM density maps and refined using several softwares.^7^ Recent works mainly focusing on the RNA dependent RNA polymerase (RdRp) NSP12,^7,8^ which is expressed in the polyprotein sequence just before the helicase, also yielded structures of the replication machinery, including low resolution cryo-EM images of the helicase. Unfortunately, the level of resolution is too low in this structure to model the ATP pocket in a catalytically competent conformation.

### Helicase Structures and Models

The recent July 2020 SARS-CoV-2 helicase structure (PDB ID 6zsl) and the almost identical SARS-CoV helicase structure from 2019 (PDB ID 6jyt)^9^ were both resolved as crystallographic dimers (Fig. 2a-b). Interestingly, the dimerization interface is different in the two cases, leading to structurally dissimilar complexes. In the cryo-EM structure of the RdRp complexed with the RNA helicase (and co-factors NSP7 and NSP8), the two helicase protomers are non-interacting (Fig. 2c). Therefore, the catalytically active form of a monomer is of interest, and a dimer may not be the biologically functioning unit.

**Figure 2.**
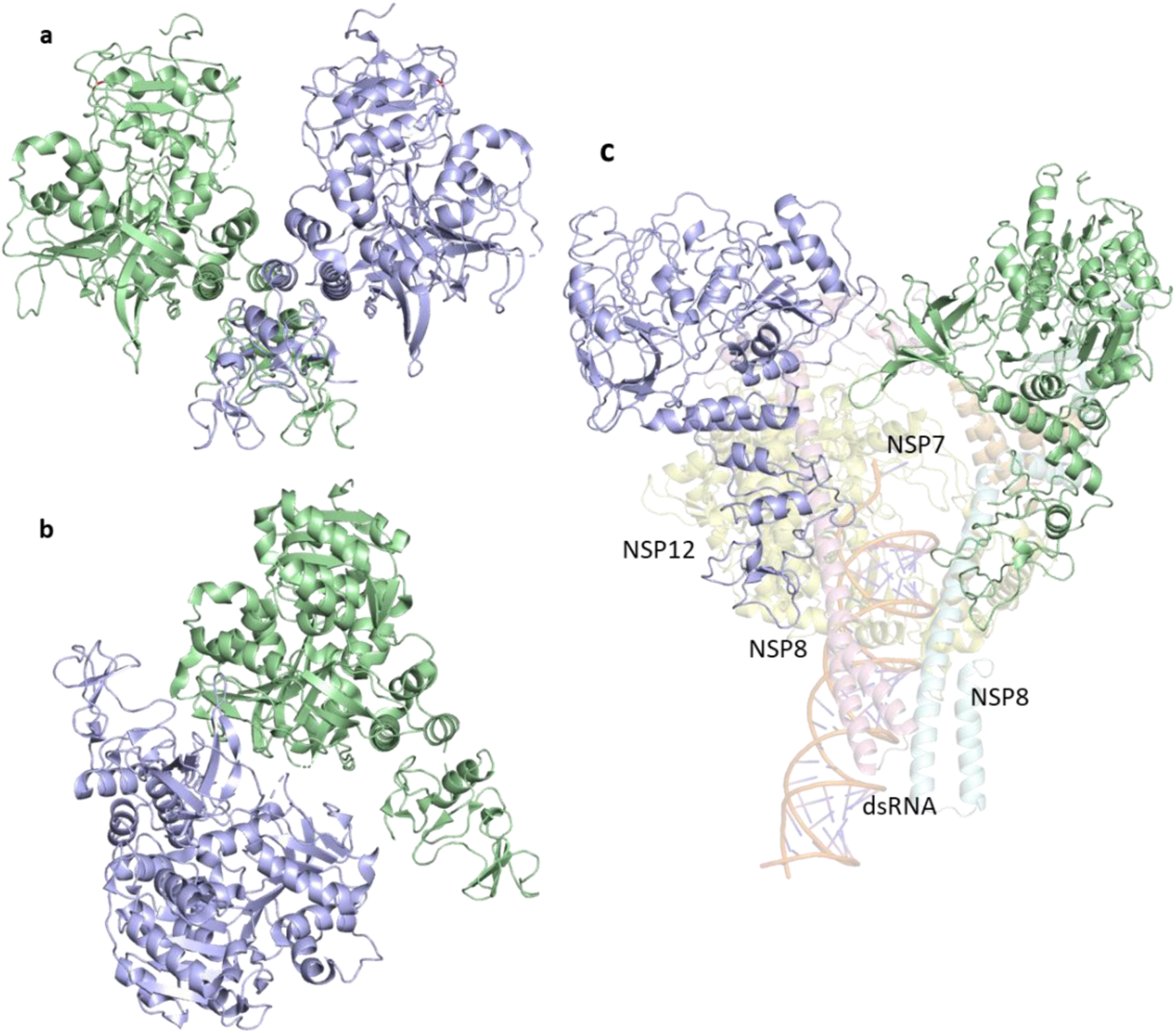
Structural comparison of the deposited PDB structures of the helicase dimer in SARS-CoV-1 (PDBID: 6jyt), SARS-CoV-2 (PDBID: 6zsl) and SARS-CoV-2 in complex with NSP7 NSP8 and NSP12 (PDBID: 6xez). The interaction between the two helicase monomers differs depending on the experimental method used to resolve the structures.

Here we present the first computational models of the SARS-CoV-2 RNA helicase with ATP and RNA substrates bound. We have performed sequence similarity searches to identify key domains and homologous sequences suggesting structurally important conserved motifs. We also performed structural alignments of available homologous helicase crystal structures to help position the bound RNA and ATP substrates (Fig. 1). A previous computational model by Hoffmann et al.^10^ described the proposed helicase ATP interactions. However, there is no RNA incorporated in these modeled structures. Here, we also present long timescale MD simulations of both the apo and ligand bound states to address the flexibility and the stability of our catalytically competent structures. Our results will help guide ongoing drug development with the identification of novel pockets.

## METHODS

### Homologous Sequence Analysis

Sequence alignments were done using BLAST with default settings and BLOSUM62 distance matrix, requesting the most similar 1000 hits from UniProtKB.^11^ Pairwise alignments for the obtained sequences were used to count identities and similarity for each residue in the SARS-CoV-2 helicase.

### Homology Modelling

Proteins with crystal structures were aligned with mustang for a combined structural-sequence alignment.^12^ The apo structure was based on PBD ID 6jyt.^9^ Missing residues and the I570V mutagenesis were constructed in pymol. The position of the ATP was determined using the coordinates of PDB ID 2xzo,^13^ as a template, modifying residues around the ATP pocket, except for Arg443 which was modeled based on PBD ID 6jim.^14^ The single stranded RNA (ssRNA) was placed based on 2xzl.^13^

### Molecular Dynamics

The helicase model was used as starting point for MD simulations. The system consists of the helicase, three Zn^2+^ ions, ATP, Mg^2+^ and ssRNA with eight uracil bases. The MD simulations were performed using NAMD 2.13,^15^ using CHARMM36 force field.^16^

The system was solvated by 50,000 – 70,000 TIP3P water molecules resulting in a box of 120 Å per side. To neutralize the system and account for a 0.15 M KCl solution we added 171 K^+^ and 189 Cl^-^ ions.^17^ Periodic boundary conditions (PBC) were used in all the simulations and the particle mesh Ewald (PME) method was used for long-range electrostatic interactions. SHAKE algorithm was deployed to constraint the covalent bonds involving hydrogen atoms. A cutoff 12 Å was used to treat non-bonding interactions.

The energy of the system was minimized using a standard protocol via steepest descent algorithm for a total number of 10,000 steps, followed by 50 ns equilibration with restrained heavy atoms (heavy atom of the backbone of the protein and the nucleic acid with an isotropic force of 1000 kJmol^-1^ nm^-1^) in constant pressure and temperature (NPT) and constant volume and temperature (NVT; up to 1ns) at 303.15 K via standard MD procedure with a time step of 2 fs.

To help equilibrate the complexes, we used a harmonic constraint on selected contacts with a force constant of 10 kcal/mol for 15 ns in our preliminary MD simulations to maintain relevant contacts. These constraints were subsequently progressively reduced and removed during the next 20 ns, using the colvar function implemented in NAMD. For the apo structure, we performed five independent unbiased MD simulations, each 1 µs long, for a total of 5 µs simulations.

To compare simulation results obtained with MD, we also carried out MD simulations using GROMACS 2018^18–21^ with the Amber ff99+ parmbsc0+chioL3 force field^22,23^ for ssRNA and Amber14SB^24^ for the helicase. To maintain the coordination of the Zn^2+^ ions, the ZAFF model was used^25^. The molecular systems were placed in a cubic box and solvated with TIP3P water molecules.^17^ The distance between the solute and the box was set to at least 14 Å. The solute was neutralized with potassium cations and then K^+^ Cl^-^ ion pairs were added to reach the salt concentration of 0.15 M. We used the ion corrections of Joung et al.^26^ as this force field has been shown to produce stable RNA structures.^27^ The parameters for Mg^2+^ are taken from Ref. 28. Long-range electrostatic interactions were treated using the particle mesh Ewald method^29,30^ with a real-space cut-off of 10 Å. The hydrogen bond lengths were restrained using P-LINCS,^19,31^ allowing a time step of 2 fs.^32^ Translational movement of the solute was removed every 1000 steps to avoid any kinetic energy build-up.^33^ After energy minimization of the solvent and equilibration of the solvated system for 10 ns using a Berendsen thermostat (τT = 1 ps) and Berendsen pressure coupling (τP = 1 ps),^32^ simulations were carried out in an NTP ensemble at a temperature of 300 K and a pressure of 1 bar using a Bussi velocity-rescaling thermostat^34^(τP = 1 ps) and a Parrinello-Rahman barostat (τP = 1 ps).^35^ During minimization and heating, all the heavy atoms of the solute were kept fixed using positional restraints. The restraints on the RNA and the protein backbone were relaxed slowly during the equilibration from 1000 kJmol^-1^ nm^2^ to 10 kJmol^-1.^ nm^2^.

Additional MD simulations, constructed using the AmberTools20 building package, were performed with the GPU version of Amber18 using the ff14SB force field to represent the protein,^36^ the ff99OL3 force field for the RNA,^37,38^ ATP parameters from Meagher et al^39^ and parameters for Mg^2+^ are taken from Ref. 28. The tetrahedral co-ordination state of the zinc was maintained using the ZAFF bonded force field.^28^ Note that additional parameters were required for the HIS-33 that interacted with the zinc via its epsilon nitrogen by reference to comparable parameters in the ZAFF using a hybrid of the center ID 4 and 6 models.^40^ For structures where ATP is bound, the octahedral co-ordination of the Mg^2+^(which involves bonds to the ATP β and γ phosphate oxygen atoms, one with oxygen of the Ser289 hydroxyl group and three structural water molecules) was constructed using the Chimera metal center builder.^41^ The solute was neutralized with potassium cations, then the protein was immersed in a box of TIP3P water molecules extending a minimum of 10 Å from the protein surface, and K^+^ Cl^-^ ion pairs were added to achieve a salt concentration of 0.14 M. MD simulations were performed in the NTP ensemble, with Berendsen temperature and pressure coupling. SHAKE was applied to all bonds involving hydrogen, allowing an MD integration timestep of 2 fs.

Long-range electrostatic interactions were treated using the particle mesh Ewald method^29,30^ with a real-space cut-off of 12 Å. To equilibrate the protein and nucleo-protein complexes, the systems was initially energy minimized with positional restraints placed upon the solute, followed by minimization of both solvent and solute. The system was then heated to 300 K in the presence of positional restraints upon the solute, which were gradually reduced from 50 kcal/mol Å^2^ to 1.0 kcal/mol Å^2^ over a timescale of 100 ps. For the apo-helicase structure, all restraints were then removed. For the ATP-RNA helicase complex which included the coordinated Mg^2+^ ion, an additional 50 ns of equilibration was performed with harmonic distance restraints (set at 2.1 Å with a spring constant of 20 kcal/mol Å^2^) to maintain the positions of coordinated atoms, and angle restraints imposing the octahedral geometry around the Mg^2+^ ion. An additional restraint was imposed to maintain the orientation of Asp374 and Glu375 to the adjacent coordinated water molecule, as observed in the MutS-ATP complex (PDB ID 1w7a,^42^). Three 1 µs simulations of the apo-structure at a salt concentration of 140 mM, and one 1.5 µs simulation in neutralizing salt were performed. We have also obtained 1 µs simulations of the ATP-helicase (two replicas), the RNA-helicase and the ATP RNA-helicase complex. For all coordinated ATP Mg^2+^ metal centers, these equilibration protocols provide stable octahedral geometries, including the complexed water molecules, during unrestrained MD over 1 µs timescales.

### Pocket analysis

For the analysis of the ATP pocket size, we used the open-source cavity detection software, Fpocket.^43^ From our MD simulations of the apo and ATP-RNA helicase, we extracted 100 equally time-spaced protein structural snapshots from the last 200 ns simulation of both systems. We performed the pocket size analysis using several parameter sets using Fpocket and compared the results for our system (see SI). We then calculated the average volume of the ATP pocket for the most optimal parameter sets and compared these between the apo and the ATP-RNA complex simulations. The parameters we modified were: −m: minimum α-sphere size; -M: maximum α-sphere size -D: first clustering α-sphere size (see SI).

## RESULTS

### Helicase domains and their sequence homology

Following previous studies, the single chain SARS-CoV-2 helicase can be divided into three domains as depicted in Fig. 1. The sequence starts with Domain 1 (residues 1-260), which features: a Zinc-binding domain (ZBD, residues 1-100), known to facilitate nucleic acid recognition; a Stalk region shaped by 2 contiguous alpha helices (residues 100-150) which functions as an interface connecting the ZBD with the rest of the Domain 1 (residues 150-260, also known as Domain 1B) that interacts with the RNA. The rest of the chain splits into RecA1 and RecA2 domains, which are well characterized in the superfamily 1B type helicases and bind ATP at their interface.^44^

We have obtained the most homologous 1000 non-redundant sequences and their alignments from the UniProtKB library. About 10% of these sequences show similarities across the whole helicase sequence and the best 95 has 400 or more positives or similarities in the sequence alignment (Fig. 3). These are all coronaviruses mainly derived from bat virome (beta and alphacoronaviruses), and affecting various hosts in the animal kingdom, including humans. Intriguingly, the next best sequence alignment only covers 235 amino acids as similar residues; all of these and subsequent aligned regions are specific to the RecA domains and range through all types of organisms.

**Figure 3.**
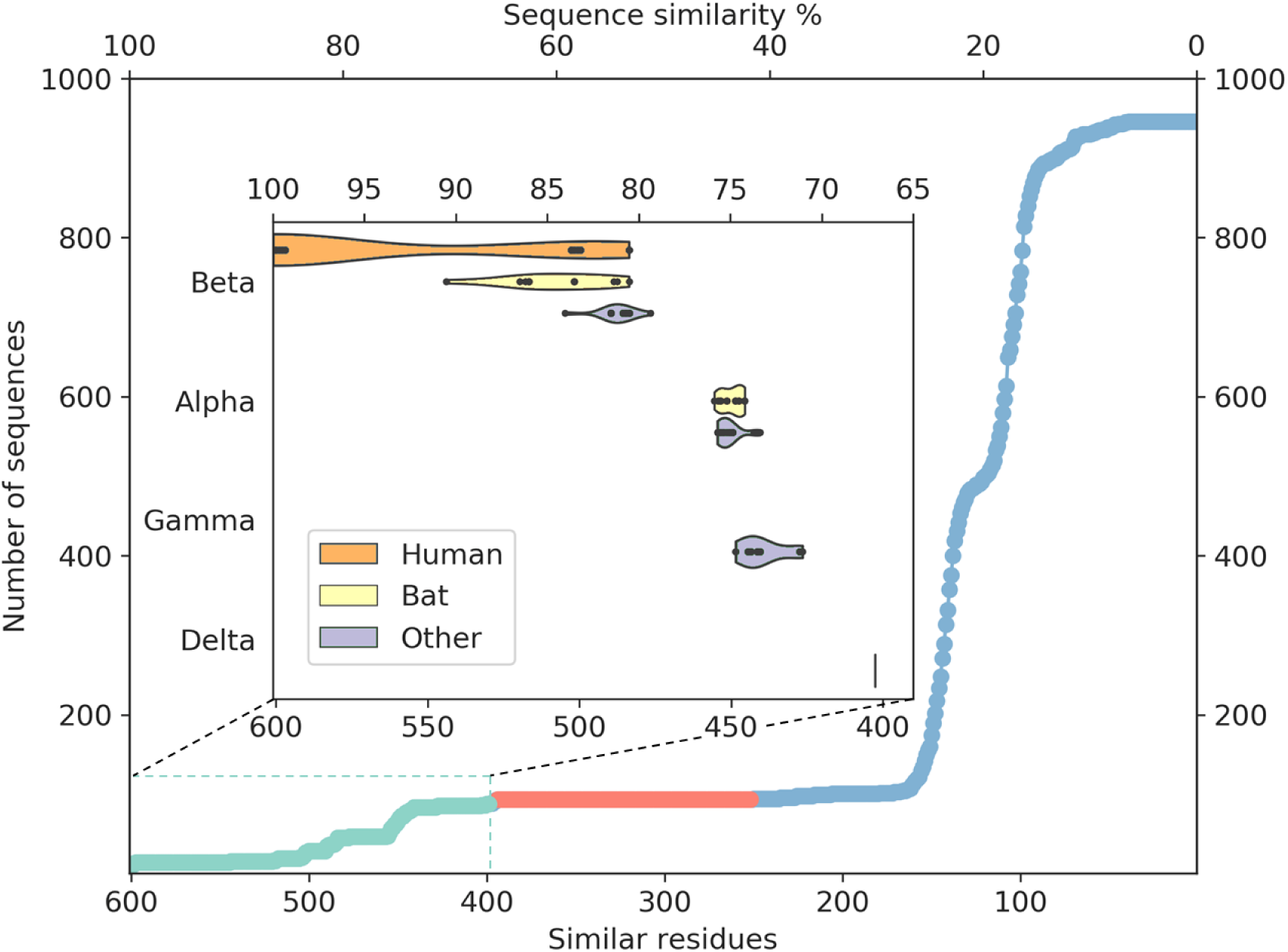
Distribution of the pairwise sequence alignments to the SARS-CoV-2 helicase. There are only members of *coronaviridae* above 398 matching residues (66%, lime circles, 95 entries). There are no sequences with medium similarity (235-394 similar residues, red circles). The closest relatives (95 sequences highlighted in lime dashed frame) are grouped in coronavirus subfamilies with principal hosts highlighted in the inset.

To evaluate any similarities to Domain 1 only, we also performed a search using only the first 230 residues. This search for sequences that match at least 70 residues resulted in the exact same 95 sequences as before, exclusively belonging to *coronaviridae*. An additional only 21 sequences match shorter segments of this domain between 1-230 residues, corresponding to a 22% sequence identity or below.

Amongst crystal structures containing ATP analogs, most helicases have a very low sequence similarity to NSP13. The closest homologues are 2xzo, 5mzn, and 6jim with 11.00%, 10.20%, and 8.40% sequence identity, respectively. Despite the low sequence identity, most residues in the ATP binding pocket are conserved. At the same time, the closest human sequence homologue based on our homology search, ZGRF1, a putative RNA helicase, shares only 22% sequence similarity, restricted to the RecA1 and RecA2 domains. This relatively narrow bandwidth of sequence similarity can be harnessed to design specific inhibitors against the coronavirus RNA helicases that do not inhibit human proteins.

### Structural model of the ATP binding site

We modelled the ATP-bound active site using the 2xzo structure as a template (Figure 7). The essential Mg^2+^ ion cofactor coordinates both the β and γ-phosphates and a conserved Ser288. The active site contains a DE of the DEAD-motif of RNA helicases. The conserved Asp374 H-bonds with the Ser288 and one of the Mg-coordinating water molecules, whereas the Glu375 is positioned as the proton acceptor.^10,45,46^ The γP is stabilized via H-bonds with Arg567 Lys289 and Gln404 through a water molecule, that are found in respectively 95, 56 and 57% of the homologous sequences analyzed, while βP forms a H-Bond with Arg443. The sugar region of the ATP likely interacts also with Glu540 and Lys320 as seen in ten and four pdb structures, respectively. Unlike the highly conserved residues recognizing the triphosphate pocket, the environment of the sugar and purine moieties (Figure 7, right) shows a greater diversity. The purine ring is stabilized through multiple π stack interactions, from one side with Arg442, while in some helicases with a tyrosine, and from the other side with His290 and Phe261. Additionally, there is a H-Bond between the amino group of the purine ring with Asn265, a residue which is more typically served by a glutamine in similar sequences.

**Figure 4.**
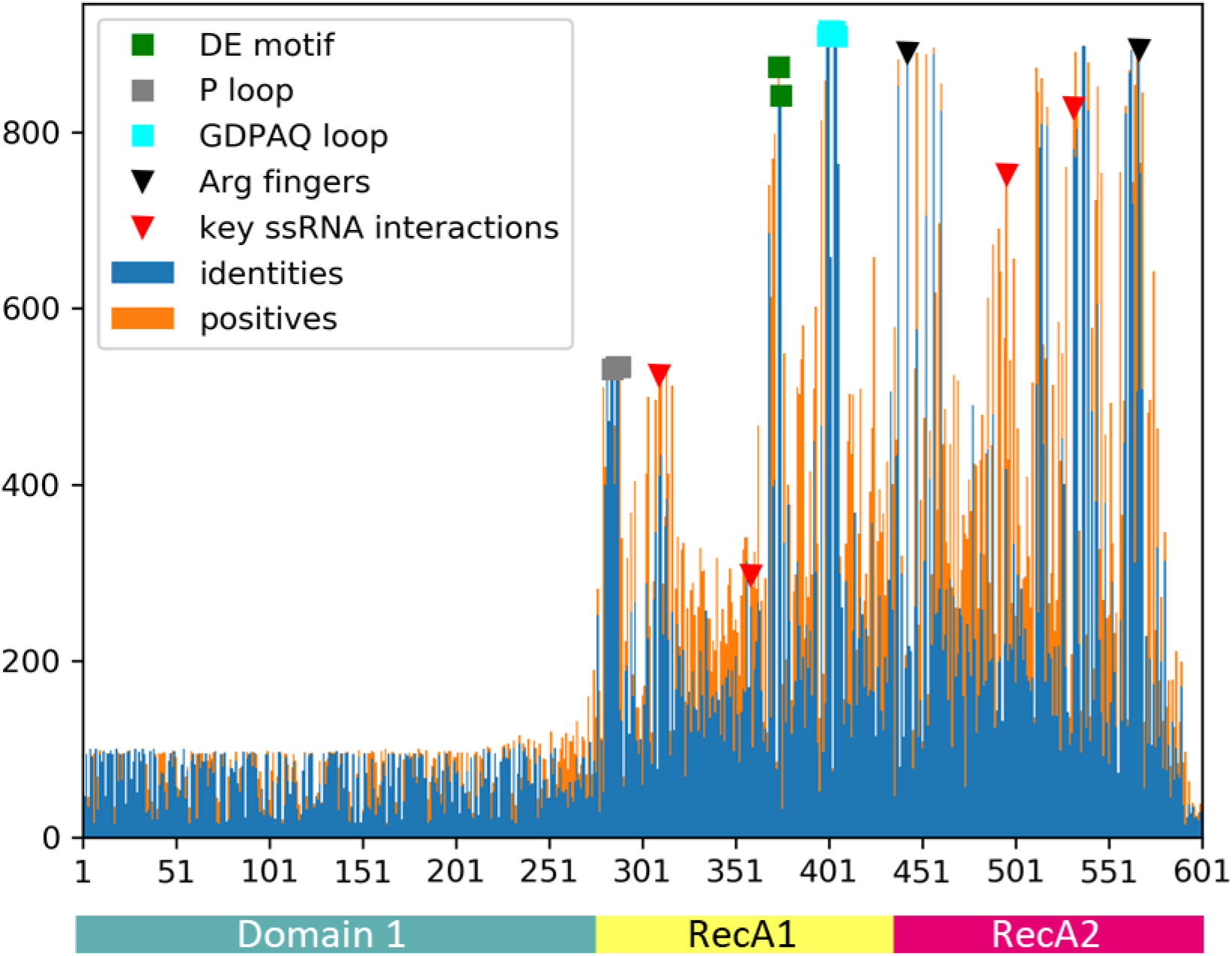
Sequence similarity (orange) and identity (blue) of the closest 946 sequences from UniProtKB using BLAST pairwise alignments to the 601-residue long SARS-CoV-2 RNA helicase. Domain 1 shows similarity only to the close relatives (95 sequences), while the RecA1 and RecA2 domains are more common across ATPase sequences. Key structural motifs are highlighted using symbols (P-loop: grey square, DE motif: green square, arginine fingers: black triangle, ssRNA interactions: red triangles).

**Figure 5.**
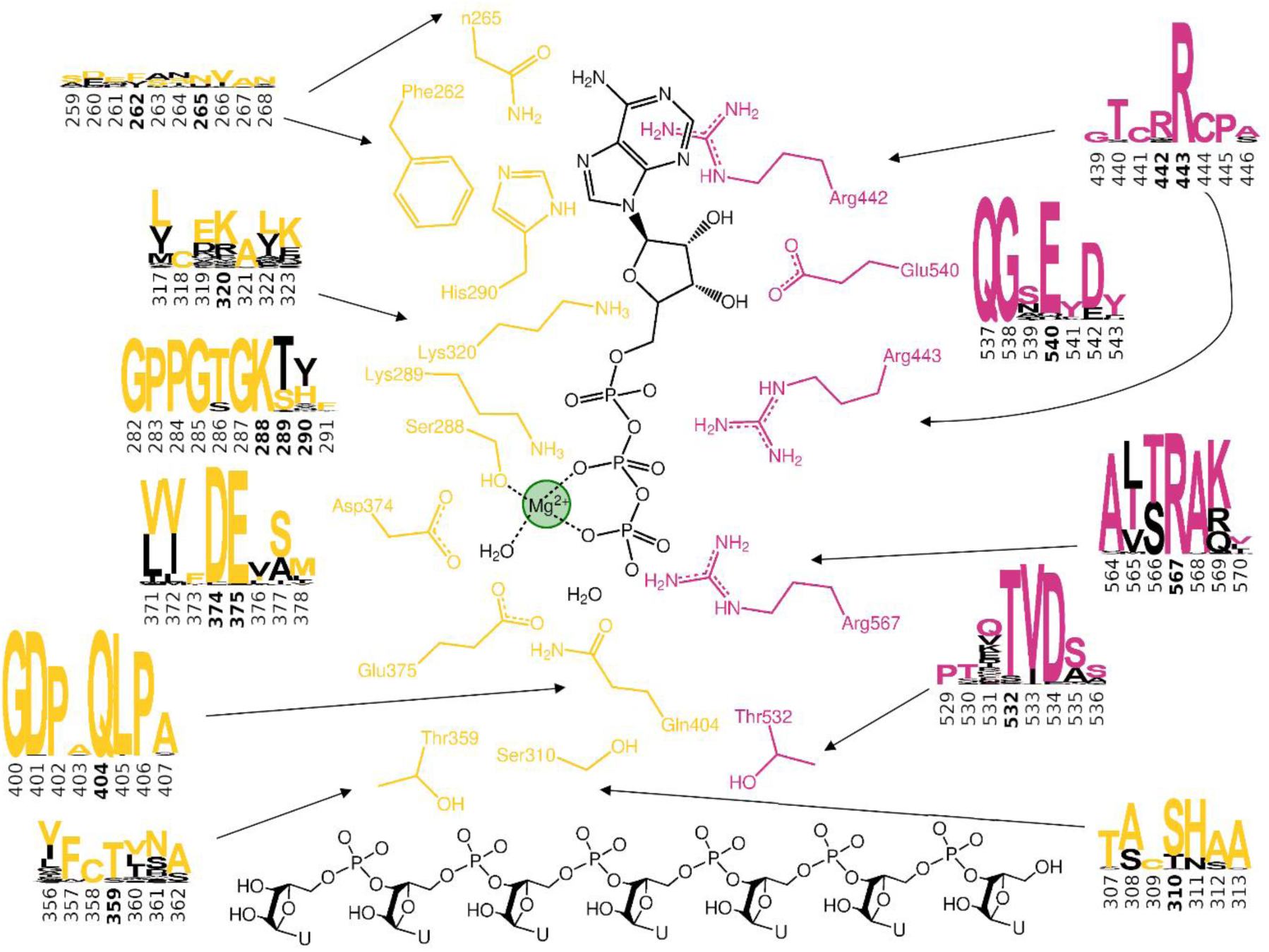
Conserved residues coordinating the ATP and RNA substrates of the SARS-CoV-2 helicase. Sequence conservation for RecA1 (orange) and RecA2 (magenta) domains are depicted in logos for each residue and its neighbors (data from Figure 4). Colored letter represent the residues in the SARS-CoV-2 helicase sequence, depicted residue indices are bold in the logos.

**Figure 6.**
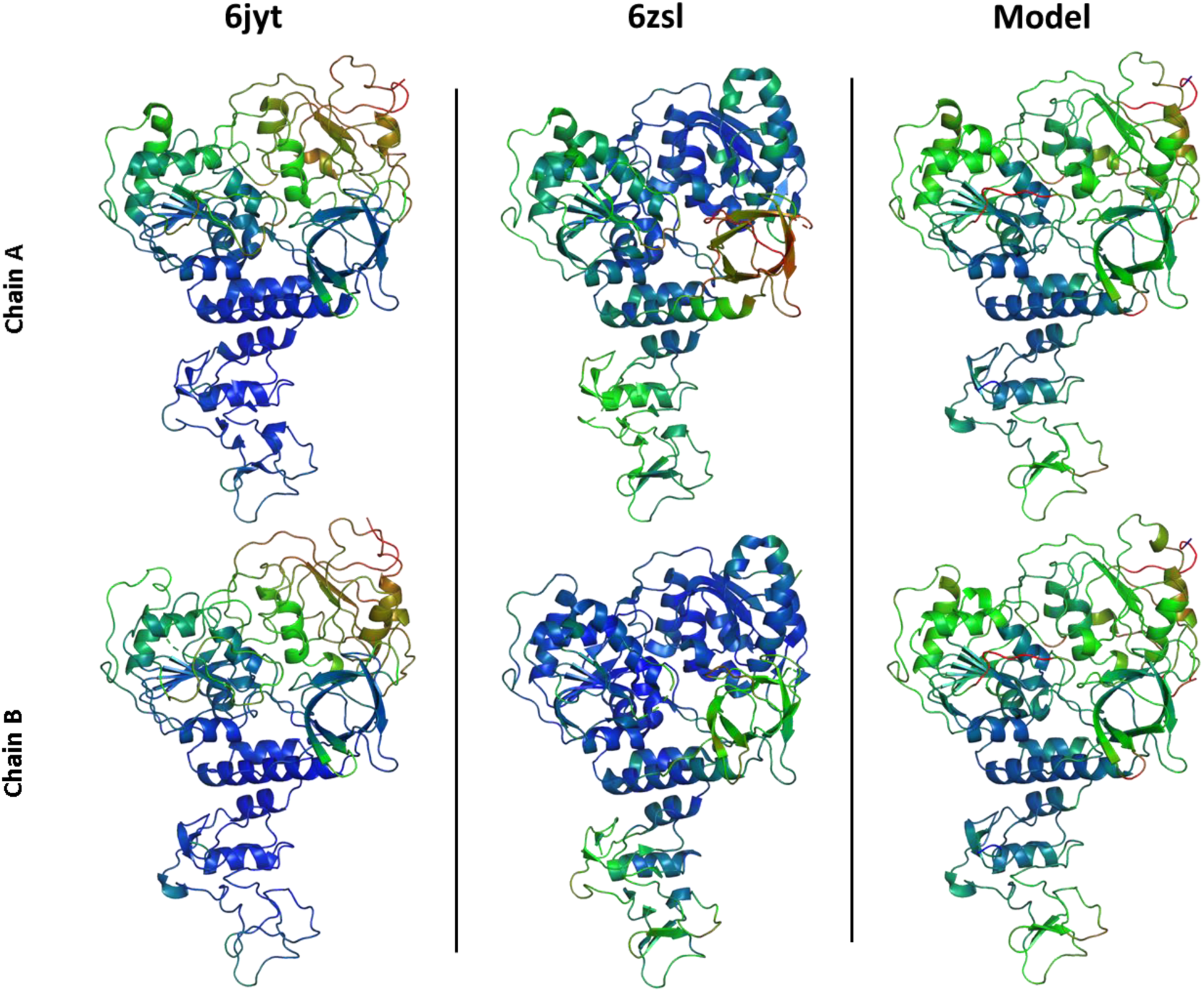
Conformational flexibility of the APO helicase protomers from 6jyt, 6zsl and our model if the apo dimer from the MD simulations. The residues are colored according to the deposited PDB B-factors (6jyt and 6zsl; from blue: low B-factor to red: high B-factor), and by the residue RMSD from the MD trajectory.

**Figure 7.**
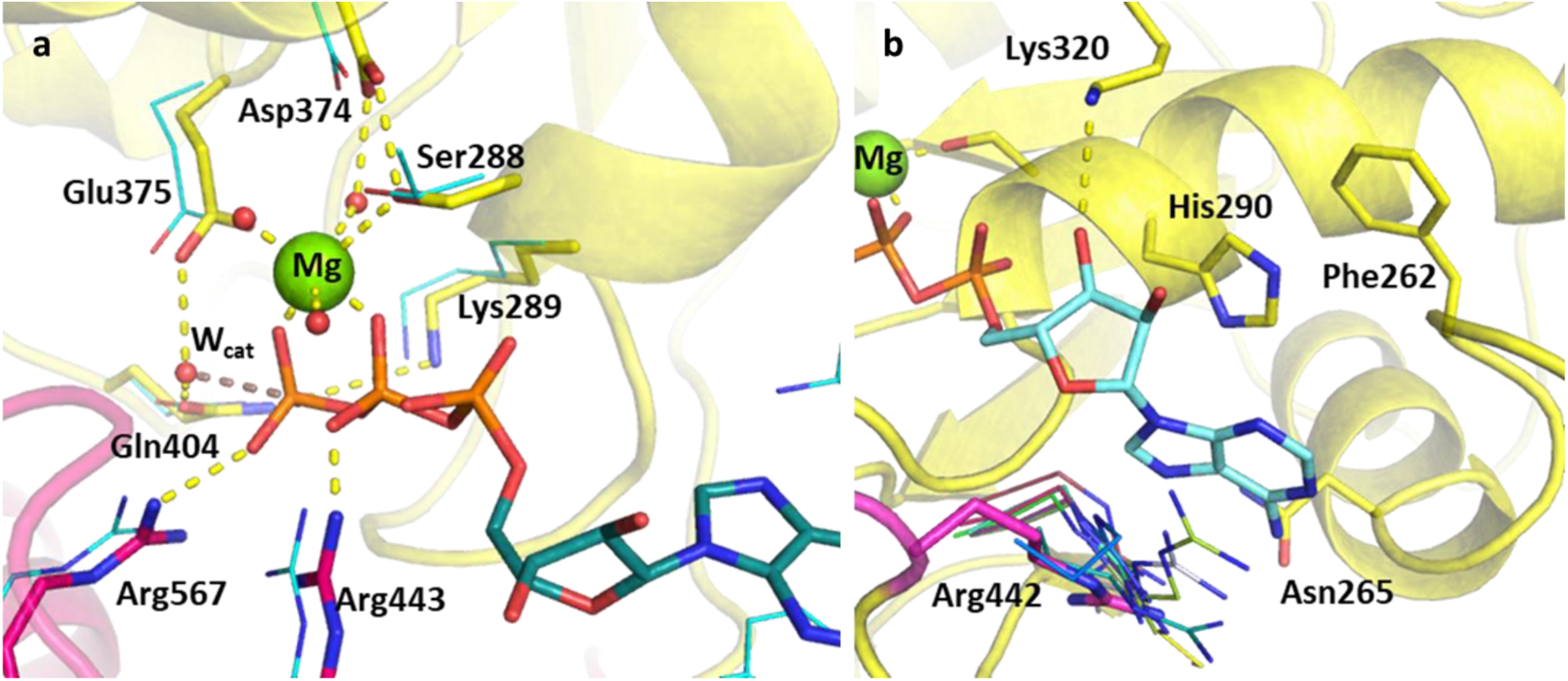
Structure of the ATP pocket aligned with homologous ATP-helicase complexes. RecA1 and RecA2 are shown in yellow and magenta, respectively. **a**) Main protein-substrate interactions of the triphosphate and magnesium ions are compared with alignment for PDB template 2xzo (cyan lines). **b**) Nucleotide binding region focusing on Arg442 (magenta sticks) is aligned with homologous arginine residues (lines, PDB structures 5k8u, 5vhc, 5xdr, 5y4z, 5y6m, 5y6n, 6adx, 6ady, 6c90 and 6jim).

A lack of specificity towards the purine group is likely due to the dual function of the SARS-CoV-2 helicase to aid the 5’-capping of the RNA by the triphosphate hydrolysis of most NTP substrates.^3^ Due to these major differences, this area of the nucleotide binding pocket can serve in the design of SARS-CoV-2-specific antiviral drugs.

### Structural model of the RNA binding site

To identify the main contact points with the ssRNA, it is first important to understand the unwinding function of the helicase. Filtering the related crystal structures to the ones containing RNA, we noticed that the RNA directionality relative to the ATP pocket is well defined (Fig. 1). The unwinding is driven by a domain-wise translocation process, which moves the RNA one base in two steps.^13,14,47^ Upon ATP hydrolysis, domain RecA2 translocate 1 base towards the 5’ end of the RNA which is then followed by the RecA1 when a new molecule of ATP binds to the enzyme.

Domain 1, being in contact with the sidechain of the RNA, does not feature specific motifs, thus allowing different RNA bases to translocaste. A long loop transitions into the RecA1 and 2 domains sandwiching the ATP pocket on the side of RNA backbone. This region, equipped with the necessary functionalities to perform the ATP hydrolysis, has a higher degree of conservation along the helicases. Both RecA domains have specific residues reliable for the contact with the RNA phosphates, depicted in Fig. 8. Thr359 in RecA1 and Thr532 are identified as the main anchoring points of the two domains. The base between these two threonine residues is coordinated by a backbone NH of His311, an interaction which is kept in RNA containing crystal structures. Ser310 is also reasonably conserved, although not directly featured in RNA coordination in this state of the enzyme.

**Figure 8.**
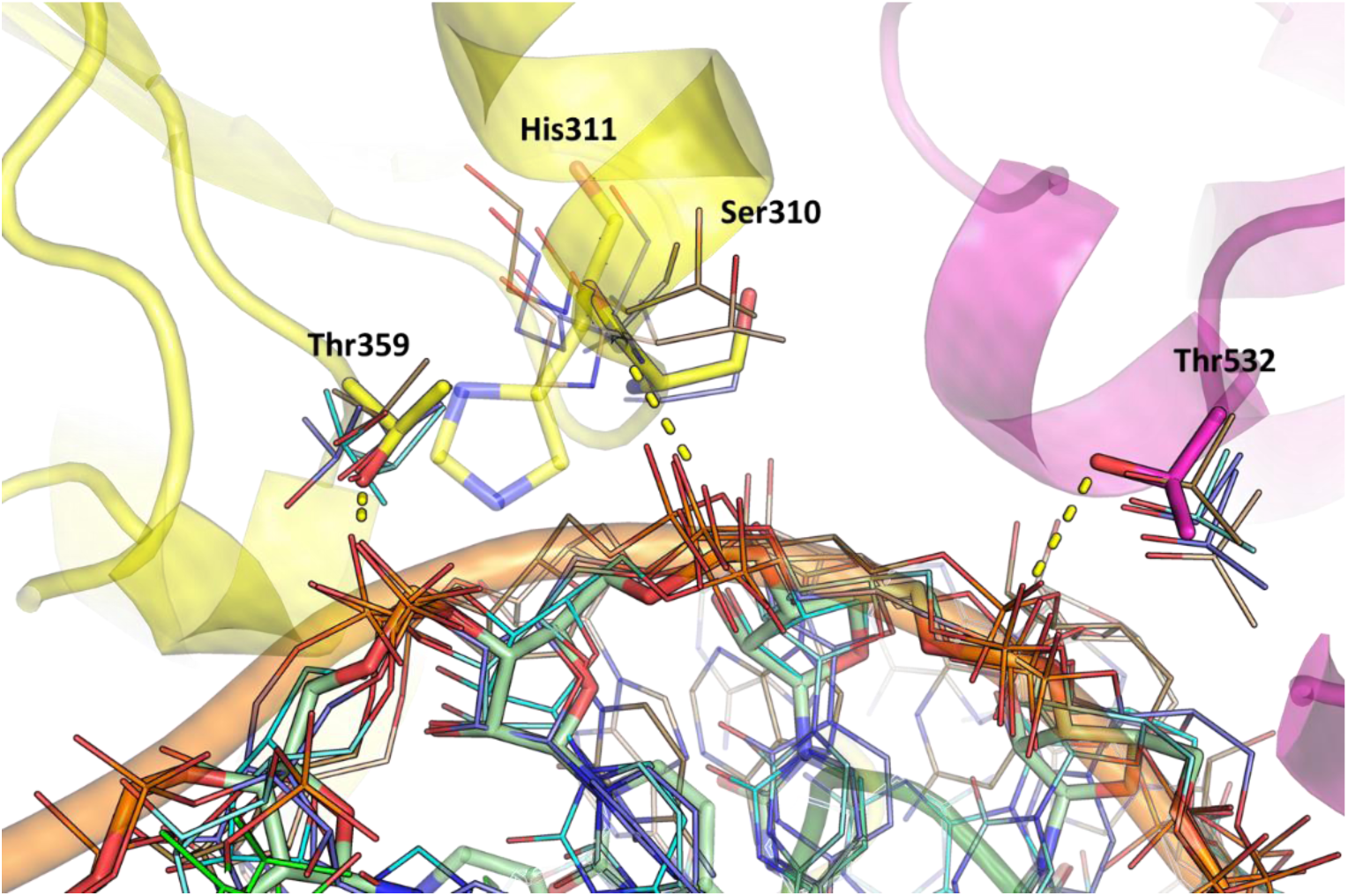
Structures of the RNA binding region aligned with existing RNA-helicase crystal structures complexed with RNA (depicted in lines). RecA1 and RecA2 domains are shown in yellow and magenta, respectively. Key residues (sticks) are labelled and H-bonds are depicted in yellow dashes.

Interestingly, the most conserved motif across the sequences is a GDP(A)Q loop interfacing between the RecA1 and RecA2 domains. This motif features Gln404, a residue which we consider to be important in the coordination of the nucleophilic water; moreover, it bridges the γP and the SH motif discussed earlier. We speculate these moieties play a role in the translocation of RecA2 upon ATP hydrolysis.

### MD simulations

#### Apo structures

All replicas of the apo structure show low flexibility and no major changes of the backbone structure of the dimer. We analyze the overall flexibility of the dimers and compare our results with the experimental b-factor obtained in 6jyt and 6zsl (Figure 6). Our model, in common with the two crystal structures, shows higher flexibility on the external shell of RecA2 domain, while the ATP and the RNA pockets appear to be more conserved. The ZBD shows low flexibility, in agreement with the b-value of 6jyt, but not with 6zsl (especially chain A), in which the temperature factor is higher, due to the different dimerization of the crystal structures.

### ATP-bound pocket

From our unbiased trajectory of the ATP-RNA-helicase complex we extract key distances along the simulations (Figure 7). The hexacoordinate Mg^2+^ shows stable bond with 3 water molecules, the OH of Ser288 and two oxygens of the β and γ phosphate of the ATP, which is essential for the preorganization of the ATP hydrolysis Further conserved contacts in the pocket are also kept stable during the simulation: including the arginine fingers, Lys288 and the DE motif which both takes part in coordinating the Mg^2+^ and the nucleophilic water. Gln404 shows a greater flexibility deep in the ATP pocket, which might suggest its role in changing the conformation during translocation. Residues participating in the adenosine coordination are less conserved and also exhibit fewer stable contacts.

### Pocket analysis

We identify the cavity size of the ATP pocket using Fpocket. We tested different combinations of parameters to find a combination that yields more robust results (see SI for details). A comparison of the average pocket volume from the apo trajectory (394.6±157.5 Å^3^) with the average pocket volume from the ATP bound trajectory (367±146 Å^3^) shows no substantial differences within the error of the analysis. This finding suggests that the ATP cavity structure is not altered significantly by the presence of ATP.

### RNA binding site

From the structural analysis and the homology modelling, we denote two important and well -conserved interactions between the RNA and the helicase. Both interactions involve a H--bond between a threonine and the O of the phosphate residue on the backbone of the RNA. Specifically, from Thr359 from RecA1 and Thr532 from RecA2. Additionally, another H-bond is made between the central RNA and the N of residue 311, this residue is not highly conserved but the interaction between the backbone of this residue and the OP of the RNA is present in several PDB structures. A key residue close to the RNA pocket is Ser310; this residue is conserved (often present as a threonine) and appears to be important for the communication between the ATP pocket and the RNA pocket.

## SUMMARY

We present the catalytically competent computational model of the SARS-CoV-2 NSP13 ATP dependent RNA helicase. Our structure is the first to host ATP and a single strand RNA with detail to the binding modes of the substrates.

The analysis of homologous sequences sheds light upon the specificity of the domain structure of the viral helicase yielding no match over 40% except close relatives from the *coronaviridae* family. Our model features two major improvements compared to the apo protein. The ATP pocket was reconstructed based on the structural conservation of the most similar crystal structures including signature motifs from phosphate binding proteins such as the DE(AD) of helicases, the P loop or the arginine fingers. Furthermore, we identified the main anchoring points of the ssRNA through the helicase, which are essential to understand the translocation driving the unwinding activity of NSP13.

With molecular dynamics, we have verified the stability of conserved interactions in our model as well as improved our initial model to host the nucleic acid. We assess the flexibility of the ATP pocket with and without the nucleotide bound detecting a well-maintained cavity.

Our work provides insight into one of the elements of the viral replication machinery, ideal for targeting by drug developers. Our structure can be the basis of structure-based compound design and screening. Moreover, elaborating the RNA translocations driven by the identified interactions can reveal other targetable states of the helicase.

## ACKNOWLEDGEMENT

This project made use of time on ARCHER granted via the UK High-End Computing Consortium for Biomolecular Simulation, HECBioSim (http://hecbiosim.ac.uk), supported by EPSRC (grant no. EP/R013012/1). We would like to thank GENCI for a generous allocation of computing time at CINES within the action “Project contre le COVID 19”. We are also grateful for resourced provided by the Leeds ARC HPC service.

