## Supporting Information for "Modelling the active SARS-CoV-2 helicase complex as a basis for structure-based inhibitor design"

Table S1. List of PDB structured used in structural comparison. Sequence similarity and RMSD is based on the mustang alignment (see methods).

| PDB ID | Sequence Length | Number of Identical Residues | RMSD | Nucleotide | NA |
|---|---|---|---|---|---|
| 2xzo | 770 | 85 | 7.2 | ADP+$AlF_4^-$ | ssRNA |
| 5mzn | 825 | 84 | 12.9 | ADP | - |
| 6jim | 668 | 56 | 10.5 | ADP+$AlF_3$ | ssRNA |
| 4f93 | 1778 | 49 | 23 | ATP | - |
| 3o8d | 823 | 44 | 20.3 | ADP | - |
| 3o8r | 822 | 41 | 20.6 | ADP+$BeF_3$ | ssRNA |
| 3rrm | 687 | 35 | 16.2 | ADP | - |
| 3i61 | 748 | 32 | 17.6 | ADP+$BeF_3$ | ssRNA |
| 4tyw | 762 | 32 | 19.8 | ADP+$BeF_3$ | ssRNA |
| 6c90 | 968 | 32 | 18.8 | ADP | - |
| 3kqu | 709 | 31 | 18.6 | ADP+$BeF_3$ | ssDNA |
| 1xtj | 689 | 30 | 13.3 | ADP | - |
| 3i62 | 750 | 29 | 17.4 | ADP+$AlF_4^-$ | ssRNA |
| 4tz0 | 753 | 29 | 18 | GDP+$BeF_3$ | ssRNA |
| 2whx | 797 | 29 | 26.2 | ADP | - |
| 5e4f | 722 | 28 | 19.5 | ADP+$AlF_4^-$ | - |
| 6uv3 | 711 | 28 | 17 | ADP+$BeF_3$ | ssRNA |
| 3kx2 | 985 | 28 | 21.9 | ADP | - |
| 6uv2 | 715 | 27 | 16.1 | ADP+$BeF_3$ | ssRNA |
| 6uv4 | 705 | 27 | 17.3 | ADP+$BeF_3$ | ssRNA |
| 5xdr | 952 | 27 | 21.3 | ADP | - |
| 5y6m | 691 | 26 | 21.1 | ADP+$AlF_3$ | - |
| 3kql | 704 | 26 | 20.2 | ADP+$AlF_3$ | ssDNA |
| 2jlr | 723 | 26 | 18.5 | ANP | - |
| 6adx | 695 | 25 | 21.3 | ADP | - |
| 6uv1 | 706 | 25 | 17.8 | ADP+$BeF_3$ | ssRNA |
| 3kqn | 711 | 25 | 18.3 | ADP+$BeF_3$ | ssDNA |
| 5k8u | 712 | 25 | 19.5 | ADP | - |
| 5vhc | 1059 | 25 | 23.2 | ADP+$BeF_3$ | - |
| 2jls | 723 | 24 | 20.2 | ADP | - |
| 5e3h | 889 | 24 | 26 | ADP+$BeF_3$ | dsRNA |
| 2jlv | 706 | 23 | 20.2 | ANP | ssRNA |
| 6ady | 695 | 23 | 21.6 | ADP | - |
| 2jlx | 712 | 23 | 19.9 | ADP+$VO_4^{3-}$ | ssRNA |
| 2pl3 | 653 | 22 | 18 | ADP | - |
| 2jlz | 721 | 22 | 18.2 | ADP | ssRNA |
| 4ljy | 738 | 22 | 27.8 | ADP | - |
| 5y6n | 704 | 21 | 19.3 | ADP | - |
| 5y4z | 714 | 21 | 20 | ANP | - |
| 3dkp | 650 | 20 | 18.9 | ADP | - |
| 3wrx | 711 | 18 | 23.3 | AGS | - |
| 3ex7 | 619 | 15 | 17.7 | ADP+$AlF_3$ | ssRNA |
| 5sup | 685 | 14 | 21.1 | ADP+$BeF_3$ | ssRNA |

Pocket analysis

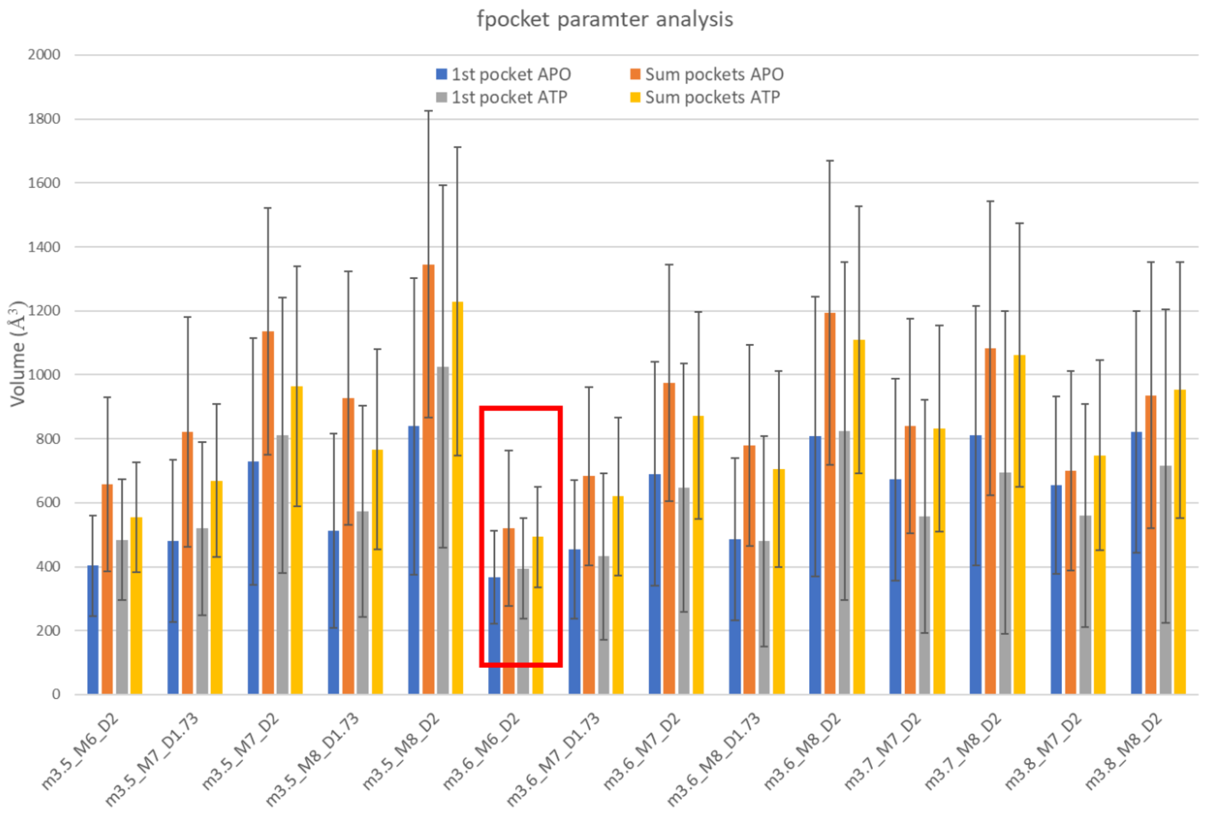


Figure S1. Statistics of ATP pocket size obtained by different parameters in Fpocket. Average volumes are displayed as columns with error bars obtained from standard deviation. Blue and grey columns represent the closest pocket to the β-phosphorus, while orange and yellow correspond to the sum of the pockets closer than 6 Å to the said atom, determined to apo simulations and one with ATP bound, respectively. Different parameters are labelled as the corresponding flags in Fpocket.
